## supplementary notes for "Powerful one-dimensional scan to detect heterotic QTL"

### Supplementary Note

In this note, we adopted all notations and equations from the subsections "MPH and the heterotic effect" and "The hQTL-ODS model" in Methods.

#### 1 The heterotic effects

In this section, we prove that the vector form of the heterotic effect (Eq. 4) is equivalent to the original definition (Eq. 3) in [1].

For any $i \in \{1, 2, \ldots, p\}$ and any hybrid individual $F$, denote by $\boldsymbol{h}_{i,F}$ the entry of $\boldsymbol{h}_i$ corresponding to $F$. Then, according to Eq. (4), $\boldsymbol{h}_{i,F}$ has the following form:

$$\begin{aligned} h_{i,F} = (\boldsymbol{T}\boldsymbol{l}_i)_F d_i + \frac{1}{2}\Bigg[ & \sum_{\substack{j=1 \\ j\neq i}}^{p} (\boldsymbol{T}(\boldsymbol{m}_i \circ \boldsymbol{m}_j))_F aa_{ij} + \sum_{\substack{j=1 \\ j\neq i}}^{p} (\boldsymbol{T}(\boldsymbol{m}_i \circ \boldsymbol{l}_j))_F ad_{ij} \\ & + \sum_{\substack{j=1 \\ j\neq i}}^{p} (\boldsymbol{T}(\boldsymbol{l}_i \circ \boldsymbol{m}_j))_F ad_{ji} + \sum_{\substack{j=1 \\ j\neq i}}^{p} (\boldsymbol{T}(\boldsymbol{l}_i \circ \boldsymbol{l}_j))_F dd_{ij} \Bigg], \end{aligned} \tag{S1}$$

where $(\boldsymbol{T}\boldsymbol{l}_i)_F$, $(\boldsymbol{T}(\boldsymbol{m}_i \circ \boldsymbol{m}_j))_F$, $(\boldsymbol{T}(\boldsymbol{m}_i \circ \boldsymbol{l}_j))_F$, $(\boldsymbol{T}(\boldsymbol{l}_i \circ \boldsymbol{m}_j))_F$ and $(\boldsymbol{T}(\boldsymbol{l}_i \circ \boldsymbol{l}_j))_F$ are the entries corresponding to $F$ in the vectors $\boldsymbol{T}\boldsymbol{l}_i$, $\boldsymbol{T}(\boldsymbol{m}_i \circ \boldsymbol{m}_j)$, $\boldsymbol{T}(\boldsymbol{m}_i \circ \boldsymbol{l}_j)$, $\boldsymbol{T}(\boldsymbol{l}_i \circ \boldsymbol{m}_j)$ and $\boldsymbol{T}(\boldsymbol{l}_i \circ \boldsymbol{l}_j)$, respectively.

Now, we need to show that Eq. (S1) is equivalent to Eq. (3). By comparing the coefficients of $d_i$, $aa_{ij}$, $ad_{ij}$, $ad_{ji}$ and $dd_{ij}$, we can see that it is sufficient to prove the following five identities:

$$(\boldsymbol{T}\boldsymbol{l}_i)_F = \begin{cases} 1, & \text{if } \ i \in R_{20} \cup R_{02} \\ 0, & \text{if } \ i \in R_{22} \cup R_{00} \end{cases} \tag{S2}$$

$$(\boldsymbol{T}(\boldsymbol{m}_i \circ \boldsymbol{m}_j))_F = \begin{cases} 1, & \text{if } \ i \in R_{20}, j \in R_{02} \ \text{ or } \ i \in R_{02}, j \in R_{20} \\ -1, & \text{if } \ i, j \in R_{20} \ \text{ or } \ i, j \in R_{02} \\ 0, & \text{if } \ i \in R_{22} \cup R_{00} \ \text{ or } \ j \in R_{22} \cup R_{00} \end{cases} \tag{S3}$$

$$(\boldsymbol{T}(\boldsymbol{m}_i \circ \boldsymbol{l}_j))_F = \begin{cases} 1, & \text{if } \ i \in R_{22}, j \in R_{20} \cup R_{02} \\ -1, & \text{if } \ i \in R_{00}, j \in R_{20} \cup R_{02} \\ 0, & \text{if } \ i \in R_{02}, \ \text{ or } \ i \in R_{20}, \ \text{ or } \ i, j \in R_{22} \cup R_{00} \end{cases} \tag{S4}$$

$$(\boldsymbol{T}(\boldsymbol{l}_i \circ \boldsymbol{m}_j))_F = \begin{cases} 1, & \text{if } \ j \in R_{22}, i \in R_{20} \cup R_{02} \\ -1, & \text{if } \ j \in R_{00}, i \in R_{20} \cup R_{02} \\ 0, & \text{if } \ j \in R_{02}, \ \text{ or } \ j \in R_{20}, \ \text{ or } \ i, j \in R_{22} \cup R_{00} \end{cases} \tag{S5}$$

$$(\boldsymbol{T}(\boldsymbol{l}_i \circ \boldsymbol{l}_j))_F = \begin{cases} 1, & \text{if } \ i, j \in R_{20} \cup R_{02} \\ 0, & \text{if } \ i \in R_{22} \cup R_{00}, \ \text{ or } \ j \in R_{22} \cup R_{00} \end{cases} \tag{S6}$$

Before proving them, we make some general observations. First, for any $(n+r)$-dimensional vector $\boldsymbol{v}$, we have $(\boldsymbol{T}\boldsymbol{v})_F = \boldsymbol{T}^F \boldsymbol{v}$, where $\boldsymbol{T}^F$ is the row vector of the matrix $\boldsymbol{T}$ that corresponds to the individual $F$. Secondly, since $\boldsymbol{T}$ is the $n \times (n+r)$ matrix transferring the vector of original trait values to that of MPH, it is clear that there are only three non-zero entries in $\boldsymbol{T}^F$ (See the example in Section 5). Namely, $\boldsymbol{T}^F_{P1} = \boldsymbol{T}^F_{P2} = -\frac{1}{2}$, and $\boldsymbol{T}^F_F = 1$, where $P1$ and $P2$ are the two parental lines of $F$, $\boldsymbol{T}^F_{P1}$, $\boldsymbol{T}^F_{P2}$ and $\boldsymbol{T}^F_F$ are the entries of $\boldsymbol{T}^F$ corresponding to $P1$, $P2$ and $F$ respectively. Let $\boldsymbol{v}^*_F = (\boldsymbol{v}_{P1}, \boldsymbol{v}_{P2}, \boldsymbol{v}_F)'$ be the three-dimensional sub-vector of $\boldsymbol{v}$ consisting of entries at the positions $P1$, $P2$ and $F$, and let $\boldsymbol{T}^{F*} = (\boldsymbol{T}^F_{P1}, \boldsymbol{T}^F_{P2}, \boldsymbol{T}^F_F) = (-\frac{1}{2}, -\frac{1}{2}, 1)$. Then, it is easy to see that $(\boldsymbol{T}\boldsymbol{v})_F = \boldsymbol{T}^F\boldsymbol{v} = \boldsymbol{T}^{F*}\boldsymbol{v}^*_F = (-\frac{1}{2}, -\frac{1}{2}, 1)(\boldsymbol{v}_{P1}, \boldsymbol{v}_{P2}, \boldsymbol{v}_F)' = \boldsymbol{v}_F - \frac{1}{2}(\boldsymbol{v}_{P1} + \boldsymbol{v}_{P2})$.

Now let's prove Eq. (S2). If $i \in R_{20} \cup R_{02}$, it means that the two parental lines are homozygous for different alleles. Then, $F$ is heterozygous. Recall that $\boldsymbol{M}_D$ is the $(n + r) \times p$ matrix of markers coded as 0 (homozygous) or 1 (heterozygous) and $\boldsymbol{l}_i$ is the $i$-th column of $\boldsymbol{M}_D$. Thus, $\boldsymbol{l}^*_i = (0, 0, 1)'$, and $(\boldsymbol{T}\boldsymbol{l}_i)_F = 1 - \frac{1}{2}(0 + 0) = 1$. If $i \in R_{22} \cup R_{00}$, then the two parental lines are homozygous for the same allele. Consequently, $F$ is also homozygous and $\boldsymbol{l}^*_i = (0, 0, 0)'$. Hence, $(\boldsymbol{T}\boldsymbol{l}_i)_F = 0 - \frac{1}{2}(0 + 0) = 0$.

Next, we prove Eq. (S3). Recall that $\boldsymbol{m}_i$ is the $i$-th column of $\boldsymbol{M}_A$ which is the $(n+r)\times p$ matrix of markers coded as 1 (homozygous for the reference allele), 0 (heterozygous) or $-1$ (homozygous for the alternative allele). If $i \in R_{20}, j \in R_{02}$, then $\boldsymbol{m}^*_i = (1, -1, 0)'$ and $\boldsymbol{m}^*_j = (-1, 1, 0)'$. If $i \in R_{02}, j \in R_{20}$, then $\boldsymbol{m}^*_i = (-1, 1, 0)'$ and $\boldsymbol{m}^*_j = (1, -1, 0)'$. In both cases, we have $(\boldsymbol{m}_i \circ \boldsymbol{m}_j)^* = (-1, -1, 0)'$. Hence, $(\boldsymbol{T}(\boldsymbol{m}_i \circ \boldsymbol{m}_j))_F = 0 - \frac{1}{2}(-1 - 1) = 1$. If $i, j \in R_{20}$, then $\boldsymbol{m}^*_i = \boldsymbol{m}^*_j = (1, -1, 0)'$. If $i, j \in R_{02}$, then $\boldsymbol{m}^*_i = \boldsymbol{m}^*_j = (-1, 1, 0)'$. In both cases, we have $(\boldsymbol{m}_i \circ \boldsymbol{m}_j)^* = (1, 1, 0)'$. As a result, $(\boldsymbol{T}(\boldsymbol{m}_i \circ \boldsymbol{m}_j))_F = 0 - \frac{1}{2}(1 + 1) = -1$. If $i \in R_{22}$, then $\boldsymbol{m}^*_i = (1, 1, 1)'$, which implies $(\boldsymbol{m}_i \circ \boldsymbol{m}_j)^* = \boldsymbol{m}^*_j$. Hence, $(\boldsymbol{T}(\boldsymbol{m}_i \circ \boldsymbol{m}_j))_F = \boldsymbol{T}^{F*}\boldsymbol{m}^*_j = \boldsymbol{m}^*_{j,F} - \frac{1}{2}(\boldsymbol{m}^*_{j,P1} + \boldsymbol{m}^*_{j,P2}) = 0$. The last equality holds true because the additive coding of any hybrid at any marker is the average of the codings of its parents. If $i \in R_{00}$, then $\boldsymbol{m}^*_i = (-1, -1, -1)'$, which implies $(\boldsymbol{m}_i \circ \boldsymbol{m}_j)^* = -\boldsymbol{m}^*_j$. Then, $(\boldsymbol{T}(\boldsymbol{m}_i \circ \boldsymbol{m}_j))_F = -\boldsymbol{T}^{F*}\boldsymbol{m}^*_j = -\boldsymbol{m}^*_{j,F} + \frac{1}{2}(\boldsymbol{m}^*_{j,P1} + \boldsymbol{m}^*_{j,P2}) = 0$. Similarly, we can derive that $(\boldsymbol{T}(\boldsymbol{m}_i \circ \boldsymbol{m}_j))_F = 0$ when $j \in R_{00}$ or $j \in R_{22}$.

Eq. (S4) and (S5) are symmetric with respect to the indices $i$ and $j$. Namely, Eq. (S5) can be obtained by exchanging $i$ and $j$ in Eq. (S4). Thus, we only need to prove Eq. (S4). If $i \in R_{22}$, we know that $\boldsymbol{m}^*_i = (1, 1, 1)'$, which implies $(\boldsymbol{m}_i \circ \boldsymbol{l}_j)^* = \boldsymbol{l}^*_j$. When $j \in R_{02} \cup R_{20}$, we have $\boldsymbol{l}^*_j = (0, 0, 1)'$. Thus, $(\boldsymbol{m}_i \circ \boldsymbol{l}_j)^* = (0, 0, 1)'$, and $((\boldsymbol{T}(\boldsymbol{m}_i \circ \boldsymbol{l}_j))_F = 1 - \frac{1}{2}(0 + 0) = 1$. When $j \in R_{22} \cup R_{00}$, we have $\boldsymbol{l}^*_j = (0, 0, 0)'$. Then, $(\boldsymbol{m}_i \circ \boldsymbol{l}_j)^* = (0, 0, 0)'$, which implies $((\boldsymbol{T}(\boldsymbol{m}_i \circ \boldsymbol{l}_j))_F = 0$. If $i \in R_{00}$, we have $\boldsymbol{m}^*_i = (-1, -1, -1)'$, which implies $(\boldsymbol{m}_i \circ \boldsymbol{l}_j)^* = -\boldsymbol{l}^*_j$. Therefore, when $j \in R_{02} \cup R_{20}$, we have $((\boldsymbol{T}(\boldsymbol{m}_i \circ \boldsymbol{l}_j))_F = -1 + \frac{1}{2}(0+0) = -1$, and when $j \in R_{22} \cup R_{00}$, we have $((\boldsymbol{T}(\boldsymbol{m}_i \circ \boldsymbol{l}_j))_F = -0 + \frac{1}{2}(0+0) = 0$. If $i \in R_{02}$ or $i \in R_{20}$, then $\boldsymbol{m}^*_i = (-1, 1, 0)'$ or $(1, -1, 0)'$. Since we know that $\boldsymbol{l}^*_j = (0, 0, 1)'$ (when $j \in R_{02} \cup R_{20}$) or $(0, 0, 0)'$ (when $j \in R_{22} \cup R_{00}$). In any case, it leads to $(\boldsymbol{m}_i \circ \boldsymbol{l}_j)^* = (0, 0, 0)'$. As a result, $((\boldsymbol{T}(\boldsymbol{m}_i \circ \boldsymbol{l}_j))_F = 0$.

Finally, we prove Eq. (S6). If $i, j \in R_{20} \cup R_{02}$, we have $\boldsymbol{l}^*_i = \boldsymbol{l}^*_j = (0, 0, 1)'$. Hence, $(\boldsymbol{l}_i \circ \boldsymbol{l}_j)^* = (0, 0, 1)'$ and $((\boldsymbol{T}(\boldsymbol{l}_i \circ \boldsymbol{l}_j))_F = 1 - \frac{1}{2}(0 + 0) = 1$. If $j \in R_{22} \cup R_{00}$, we have $\boldsymbol{l}^*_j = (0, 0, 0)'$. If $i \in R_{22} \cup R_{00}$, we have $\boldsymbol{l}^*_i = (0, 0, 0)'$. In both cases, $(\boldsymbol{l}_i \circ \boldsymbol{l}_j)^* = (0, 0, 0)'$, which implies $((\boldsymbol{T}(\boldsymbol{l}_i \circ \boldsymbol{l}_j))_F = 0$.

### 2 Dealing with non-independent residuals

Recall the null model of hQTL-ODS:

$$\boldsymbol{y}_{MPH} = \boldsymbol{X}\boldsymbol{\alpha} + \boldsymbol{g}_D + \boldsymbol{g}_{AA} + \boldsymbol{g}_{AD} + \boldsymbol{g}_{DD} + \boldsymbol{\varepsilon}, \tag{S7}$$

which is the same as Eq. (7). The model assumptions are $\boldsymbol{g}_D \sim N(0, \boldsymbol{K}_D\sigma^2_D)$, $\boldsymbol{g}_{AA} \sim N(0, \boldsymbol{K}_{AA}\sigma^2_{AA})$, $\boldsymbol{g}_{AD} \sim N(0, \boldsymbol{K}_{AD}\sigma^2_{AD})$, $\boldsymbol{g}_{DD} \sim N(0, \boldsymbol{K}_{DD}\sigma^2_{DD})$, $\boldsymbol{\varepsilon} \sim N(0, \boldsymbol{T}\boldsymbol{T}'\sigma^2_\varepsilon)$ and $\boldsymbol{\alpha}$ is a vector of fixed effects. This is a linear mixed model with multiple random terms and non-independent residuals. Since most software packages do not provide the option of user-specified covariance matrix for residuals, we applied a linear transformation to the model so that the residuals become independent.

More precisely, let $\boldsymbol{T}\boldsymbol{T}' = \boldsymbol{U}\boldsymbol{\Lambda}\boldsymbol{U}'$ be the eigen-decomposition of $\boldsymbol{T}\boldsymbol{T}'$. Thus, $\boldsymbol{U}$ is an orthogonal matrix and $\boldsymbol{\Lambda}$ is a diagonal matrix with positive diagonal elements (because $\boldsymbol{T}\boldsymbol{T}'$ is positive-definite).

We can write $\boldsymbol{\Lambda} = \mathrm{diag}(\lambda_1, \lambda_2, \ldots, \lambda_n)$, where $\lambda_i > 0$ for any $i \in \{1, 2, \ldots, p\}$. Define the square toot of $\boldsymbol{\Lambda}$ as $\boldsymbol{\Lambda}^{\frac{1}{2}} = \mathrm{diag}(\sqrt{\lambda_1}, \sqrt{\lambda_2}, \ldots, \sqrt{\lambda_n})$.

Now, let $\boldsymbol{V} = \boldsymbol{U}\boldsymbol{\Lambda}^{\frac{1}{2}}$ and left-multiply both sides of Eq. (S7) with $\boldsymbol{V}^{-1} = \boldsymbol{\Lambda}^{-\frac{1}{2}}\boldsymbol{U}'$, we obtain

$$\boldsymbol{V}^{-1}\boldsymbol{y}_{MPH} = \boldsymbol{V}^{-1}\boldsymbol{X}\boldsymbol{\alpha} + \boldsymbol{V}^{-1}\boldsymbol{g}_D + \boldsymbol{V}^{-1}\boldsymbol{g}_{AA} + \boldsymbol{V}^{-1}\boldsymbol{g}_{AD} + \boldsymbol{V}^{-1}\boldsymbol{g}_{DD} + \boldsymbol{V}^{-1}\boldsymbol{\varepsilon}. \tag{S8}$$

Then, we can derive that the residuals after transformation are independent. Namely,

$$\begin{aligned}\mathrm{var}(\boldsymbol{V}^{-1}\boldsymbol{\varepsilon}) &= \boldsymbol{V}^{-1}\,\mathrm{var}(\boldsymbol{\varepsilon})\boldsymbol{V} = \boldsymbol{V}^{-1}(\boldsymbol{T}\boldsymbol{T}')\boldsymbol{V}\sigma^2_\varepsilon \\ &= \boldsymbol{\Lambda}^{-\frac{1}{2}}\boldsymbol{U}'(\boldsymbol{U}\boldsymbol{\Lambda}\boldsymbol{U}')\boldsymbol{U}\boldsymbol{\Lambda}^{-\frac{1}{2}}\sigma^2_\varepsilon = \boldsymbol{\Lambda}^{-\frac{1}{2}}\boldsymbol{\Lambda}\boldsymbol{\Lambda}^{-\frac{1}{2}}\sigma^2_\varepsilon \\ &= \boldsymbol{I}_n\sigma^2_\varepsilon\end{aligned}$$

Thus, the transformed model (S8) can be solved by using standard software packages which require independent residuals. Note that one could also use the Chelosky decomposition instead of the spectral decomposition to find the transformation matrix.

### 3 Fixing the ratio of variance components

Recall the alternative model of hQTL-ODS:

$$\boldsymbol{y}_{MPH} = \boldsymbol{X}\boldsymbol{\alpha} + \boldsymbol{h}_i + \boldsymbol{g}_D + \boldsymbol{g}_{AA} + \boldsymbol{g}_{AD} + \boldsymbol{g}_{DD} + \boldsymbol{\varepsilon}, \tag{S9}$$

which is the same as Eq. (8). The model assumptions are the same as for Eq. (S7), and in addition, $\boldsymbol{h}_i \sim N(0, \boldsymbol{H}\sigma^2_h)$.

This model has to be solved for all $i \in \{1, 2, \ldots, p\}$ and is time-consuming. To accelerate the procedure, we apply an approximation that is similar to the so-called "population parameters previously determined" (P3D) approach [2].

More precisely, denote by $\hat{\sigma}^2_D$, $\hat{\sigma}^2_{AA}$, $\hat{\sigma}^2_{AD}$, $\hat{\sigma}^2_{DD}$ and $\hat{\sigma}^2_\varepsilon$ the estimates of the corresponding variance components in the null model (Eq. S7), and let $\rho_D = \hat{\sigma}^2_D/\hat{\sigma}^2_\varepsilon$, $\rho_{AA} = \hat{\sigma}^2_{AA}/\hat{\sigma}^2_\varepsilon$, $\rho_{AD} = \hat{\sigma}^2_{AD}/\hat{\sigma}^2_\varepsilon$, $\rho_{DD} = \hat{\sigma}^2_{DD}/\hat{\sigma}^2_\varepsilon$. Then, we consider the following simplified model:

$$\boldsymbol{y}_{MPH} = \boldsymbol{X}\boldsymbol{\alpha} + \boldsymbol{h}_i + \boldsymbol{g}, \tag{S10}$$

where $\boldsymbol{y}_{MPH}$, $\boldsymbol{X}\boldsymbol{\alpha}$ and $\boldsymbol{h}_i$ are the same as in Eq. (S9), $\boldsymbol{g} \sim N(0, \boldsymbol{K}\sigma^2_\varepsilon)$, and $\boldsymbol{K} = \rho_D\boldsymbol{K}_D + \rho_{AA}\boldsymbol{K}_{AA} + \rho_{AD}\boldsymbol{K}_{AD} + \rho_{DD}\boldsymbol{K}_{DD} + \boldsymbol{I}_n$. Model (S10) can be treated as an approximation of (S9) by fixing the ratios of the variance components $\sigma^2_D$, $\sigma^2_{AA}$, $\sigma^2_{AD}$ and $\sigma^2_{DD}$ to the residual variance $\sigma^2_\varepsilon$. Compared with (S9), which has six random vectors with different covariance matrices (including the residuals), model (S10) has only two random terms. In this way, the computational efficiency is improved. Of course, here the covariance matrix $\boldsymbol{K}$ is not an identity matrix and similar techniques as in Section 2 can be applied to transform (S10) into a model with independent residuals.

### 4 Efficiently calculating the covariance matrices

In this section, we discuss how to efficiently calculate the covariance matrices $\boldsymbol{K}_{AA}$, $\boldsymbol{K}_{AD}$, $\boldsymbol{K}_{DD}$ and $\boldsymbol{H}_i$ for all $i \in \{1, 2, \ldots, p\}$.

#### 4.1 Calculating $\boldsymbol{K}_{AA}$, $\boldsymbol{K}_{AD}$ and $\boldsymbol{K}_{DD}$

First, we take $\boldsymbol{K}_{AA}$ as an example. Recall that $\boldsymbol{K}_{AA} = \boldsymbol{T}\boldsymbol{M}_{AA}\boldsymbol{M}'_{AA}\boldsymbol{T}'/c_{AA}$, where $c_{AA} = \mathrm{tr}(\boldsymbol{T}\boldsymbol{M}_{AA}\boldsymbol{M}'_{AA}\boldsymbol{T}')$, $\boldsymbol{M}_{AA}$ is an $(n+r) \times p(p-1)/2$ matrix whose columns consist of $\boldsymbol{m}_i \circ \boldsymbol{m}_j$ for all $i, j$ such that $1 \le i < j \le p$. If we calculate $\boldsymbol{K}_{AA}$ directly using the formula $\boldsymbol{K}_{AA} = \boldsymbol{T}\boldsymbol{M}_{AA}\boldsymbol{M}'_{AA}\boldsymbol{T}'/c_{AA}$, the time complexity is $\mathcal{O}(n^2p^2)$ due to calculating the matrix product $\boldsymbol{M}_{AA}\boldsymbol{M}'_{AA}$. In [3], an efficient way of calculating epistatic genomic relationship matrices was developed. In particular, the matrix product $\boldsymbol{M}_{AA}\boldsymbol{M}'_{AA}$ can be calculated using the following formula (see also [4, 5]):

$$\boldsymbol{M}_{AA}\boldsymbol{M}'_{AA} = \frac{1}{2}(\boldsymbol{M}_A\boldsymbol{M}'_A) \circ (\boldsymbol{M}_A\boldsymbol{M}'_A) - \frac{1}{2}(\boldsymbol{M}_A \circ \boldsymbol{M}_A)(\boldsymbol{M}_A \circ \boldsymbol{M}_A)' \tag{S11}$$

Using the above formula, we do not have to explicitly generate the large matrix $\boldsymbol{M}_{AA}$ and the time complexity for calculating $\boldsymbol{M}_{AA}\boldsymbol{M}'_{AA}$ is reduced to $\mathcal{O}(n^2p)$. To see this, note that the Hadarmad product $\boldsymbol{M}_A \circ \boldsymbol{M}_A$ only requires $\mathcal{O}(np)$ time, the matrix products $\boldsymbol{M}_A\boldsymbol{M}'_A$ and $(\boldsymbol{M}_A \circ \boldsymbol{M}_A)(\boldsymbol{M}_A \circ \boldsymbol{M}_A)'$ require $\mathcal{O}(n^2p)$ time, and the Hadamard product $(\boldsymbol{M}_A \circ \boldsymbol{M}_A)(\boldsymbol{M}_A \circ \boldsymbol{M}_A)'$ takes $\mathcal{O}(n^2)$ time.

Similarly, to calculate the matrices $\boldsymbol{K}_{AD} = \boldsymbol{T}\boldsymbol{M}_{AD}\boldsymbol{M}'_{AD}\boldsymbol{T}'/c_{AD}$ and $\boldsymbol{K}_{DD} = \boldsymbol{T}\boldsymbol{M}_{DD}\boldsymbol{M}'_{DD}\boldsymbol{T}'/c_{DD}$, we do not explicitly generate $\boldsymbol{M}_{AD}$ and $\boldsymbol{M}_{DD}$. Instead, we use the following formulas [3]:

$$\begin{aligned}\boldsymbol{M}_{AD}\boldsymbol{M}'_{AD} &= (\boldsymbol{M}_A\boldsymbol{M}'_A) \circ (\boldsymbol{M}_D\boldsymbol{M}'_D) - (\boldsymbol{M}_A \circ \boldsymbol{M}_D)(\boldsymbol{M}_A \circ \boldsymbol{M}_D)' \\ \boldsymbol{M}_{DD}\boldsymbol{M}'_{DD} &= \frac{1}{2}(\boldsymbol{M}_D\boldsymbol{M}'_D) \circ (\boldsymbol{M}_D\boldsymbol{M}'_D) - \frac{1}{2}(\boldsymbol{M}_D \circ \boldsymbol{M}_D)(\boldsymbol{M}_D \circ \boldsymbol{M}_D)'\end{aligned} \tag{S12}$$

#### 4.2 Calculating $\boldsymbol{H}_i$

Recall that $\boldsymbol{H}_i$ is defined as $\boldsymbol{Z}_i\boldsymbol{Z}'_i/c_i$, where $c_i = \mathrm{tr}(\boldsymbol{Z}_i\boldsymbol{Z}'_i)$, $\boldsymbol{Z}_i$ is the $n \times (4p-3)$ matrix whose columns consist of $\boldsymbol{T}\boldsymbol{l}_i$, $\boldsymbol{T}(\boldsymbol{m}_i \circ \boldsymbol{m}_j)$, $\boldsymbol{T}(\boldsymbol{m}_i \circ \boldsymbol{l}_j)$, $\boldsymbol{T}(\boldsymbol{l}_i \circ \boldsymbol{m}_j)$ and $\boldsymbol{T}(\boldsymbol{l}_i \circ \boldsymbol{l}_j)$, for all $j \in \{1, 2, \ldots, p\}$ and $j \neq i$. If we generate $\boldsymbol{Z}_i$ and calculate the matrix product $\boldsymbol{Z}_i\boldsymbol{Z}'_i$, the time complexity is $\mathcal{O}(n^2p)$. Since we need to calculate $\boldsymbol{H}_i$ for all $i \in \{1, 2, \ldots, p\}$, the total complexity is $\mathcal{O}(n^2p^2)$. In the following, we show that it is not necessary to explicitly generate $\boldsymbol{Z}_i$.

Let $\boldsymbol{M}^i_{AA}$ be the $(n+r) \times (p-1)$ matrix whose columns are $\boldsymbol{m}_i \circ \boldsymbol{m}_j$ for all $j \in \{1, 2, \ldots, p\}$ and $j \neq i$. Similarly, define $\boldsymbol{M}^i_{AD}$, $\boldsymbol{M}^i_{DA}$ and $\boldsymbol{M}^i_{DD}$ be the matrices whose columns consist of $\boldsymbol{m}_i \circ \boldsymbol{l}_j$, $\boldsymbol{l}_i \circ \boldsymbol{m}_j$ and $\boldsymbol{l}_i \circ \boldsymbol{l}_j$, respectively. Then, by the definition of $\boldsymbol{Z}_i$, we have

$$\begin{aligned}\boldsymbol{Z}_i\boldsymbol{Z}'_i = \boldsymbol{T}\boldsymbol{l}_i\boldsymbol{l}'_i\boldsymbol{T}' + \boldsymbol{T}\boldsymbol{M}^i_{AA}(\boldsymbol{M}^i_{AA})'\boldsymbol{T}' + \boldsymbol{T}\boldsymbol{M}^i_{AD}(\boldsymbol{M}^i_{AD})'\boldsymbol{T}' \\ + \boldsymbol{T}\boldsymbol{M}^i_{DA}(\boldsymbol{M}^i_{DA})'\boldsymbol{T}' + \boldsymbol{T}\boldsymbol{M}^i_{DD}(\boldsymbol{M}^i_{DD})'\boldsymbol{T}'.\end{aligned} \tag{S13}$$

The time complexity of calculating $\boldsymbol{T}\boldsymbol{l}_i\boldsymbol{l}'_i\boldsymbol{T}'$ is $\mathcal{O}(n^2(n+r))$ and it takes $\mathcal{O}(n^2(n+r)p)$ time for all $i \in \{1, 2, \ldots, p\}$. Since $p$ is usually much larger than $n+r$, it is expected that $n^2(n+r)p$ is much smaller than $n^2p^2$. Thus, the calculation of this term is not the bottleneck. However, each of the remaining terms in (S13) requires $\mathcal{O}(n(n+r)p)$ time. Thus, the total complexity for all $i$ is $\mathcal{O}(n(n+r)p^2)$ and we need to develop an efficient way for calculating these terms.

Let's take $\boldsymbol{M}^i_{AA}(\boldsymbol{M}^i_{AA})'$ as an example. For any $s, t$ such that $1 \leq s \leq n+r, 1 \leq t \leq p$, denote by $m_{st}$ the $(s,t)$-entry of $\boldsymbol{M}_A$. Because $\boldsymbol{M}^i_{AA}$ has columns $\boldsymbol{m}_i \circ \boldsymbol{m}_j$ for $j \in \{1, 2, \ldots, p\}$ and $j \neq i$, it is convenient to consider its columns indexed by $\{1, 2, \ldots, i-1, i+1, \ldots, p\}$ rather than $\{1, 2, \ldots, p-1\}$. Then, for any $s, t$ such that $1 \leq s \leq n+r, 1 \leq t \leq p$ and $t \neq i$, we know that the $(s,t)$-entry of $\boldsymbol{M}^i_{AA}$ is $m_{st}m_{si}$.

Hence, for any $k, l \in \{1, 2, \ldots, n+r\}$, the $(k,l)$-entry of $\boldsymbol{M}^i_{AA}(\boldsymbol{M}^i_{AA})'$ is calculated as following:

$$\begin{aligned}
(\boldsymbol{M}^i_{AA}(\boldsymbol{M}^i_{AA})')_{kl} &= \sum_{\substack{s=1\\ s\neq i}}^{p} (\boldsymbol{M}^i_{AA})_{ks}(\boldsymbol{M}^i_{AA})_{ls} \\
&= \sum_{\substack{s=1\\ s\neq i}}^{p} (m_{ks}m_{ki})(m_{ls}m_{li}) \\
&= m_{ki}m_{li} \cdot \sum_{\substack{s=1\\ s\neq i}}^{p} m_{ks}m_{ls} \\
&= m_{ki}m_{li} \cdot \sum_{s=1}^{p} m_{ks}m_{ls} - m_{ki}^2 m_{li}^2 \\
&= (\boldsymbol{m}_i\boldsymbol{m}_i')_{kl}(\boldsymbol{M}_A\boldsymbol{M}_A')_{kl} - ((\boldsymbol{m}_i\boldsymbol{m}_i') \circ (\boldsymbol{m}_i\boldsymbol{m}_i'))_{kl}.
\end{aligned}$$

Therefore, we have the following formula

$$\boldsymbol{M}^i_{AA}(\boldsymbol{M}^i_{AA})' = (\boldsymbol{m}_i\boldsymbol{m}_i') \circ (\boldsymbol{M}_A\boldsymbol{M}_A') - (\boldsymbol{m}_i\boldsymbol{m}_i') \circ (\boldsymbol{m}_i\boldsymbol{m}_i'). \tag{S14}$$

With this formula, we do not need to explicitly generate the matrix $\boldsymbol{M}^i_{AA}$. Instead, although the matrix product $\boldsymbol{M}_A\boldsymbol{M}_A'$ takes $\mathcal{O}((n+r)^2 p)$ time, it is independent of $i$. Thus, we only need to calculate it once. The other parts of calculations have to be repeated for all $i$. The product $\boldsymbol{m}_i\boldsymbol{m}_i'$ takes $\mathcal{O}((n+r)^2)$ time and the same for the Hadamard products $(\boldsymbol{l}_i\boldsymbol{l}_i') \circ (\boldsymbol{M}_A\boldsymbol{M}_A')$ and $(\boldsymbol{m}_i\boldsymbol{m}_i') \circ (\boldsymbol{m}_i\boldsymbol{m}_i')$. Once $\boldsymbol{M}^i_{AA}(\boldsymbol{M}^i_{AA})'$ is calculated, the matrix product $\boldsymbol{T}\boldsymbol{M}^i_{AA}(\boldsymbol{M}^i_{AA})'\boldsymbol{T}'$ takes $\mathcal{O}(n(n+r)^2)$ time. Thus, the total complexity of calculating $\boldsymbol{T}\boldsymbol{M}^i_{AA}(\boldsymbol{M}^i_{AA})'\boldsymbol{T}'$ for all $i$ using Eq. (S14) is $\mathcal{O}(n(n+r)^2 p)$.

Similarly, we can derive the following formulas for $\boldsymbol{M}^i_{AD}(\boldsymbol{M}^i_{AD})'$, $\boldsymbol{M}^i_{DA}(\boldsymbol{M}^i_{DA})'$ and $\boldsymbol{M}^i_{DD}(\boldsymbol{M}^i_{DD})'$:

$$\begin{aligned}
\boldsymbol{M}^i_{AD}(\boldsymbol{M}^i_{AD})' &= (\boldsymbol{m}_i\boldsymbol{l}_i') \circ (\boldsymbol{M}_A\boldsymbol{M}_D') - (\boldsymbol{m}_i\boldsymbol{l}_i') \circ (\boldsymbol{m}_i\boldsymbol{l}_i') \\
\boldsymbol{M}^i_{DA}(\boldsymbol{M}^i_{DA})' &= (\boldsymbol{l}_i\boldsymbol{m}_i') \circ (\boldsymbol{M}_D\boldsymbol{M}_A') - (\boldsymbol{l}_i\boldsymbol{m}_i') \circ (\boldsymbol{l}_i\boldsymbol{m}_i') \\
\boldsymbol{M}^i_{DD}(\boldsymbol{M}^i_{DD})' &= (\boldsymbol{l}_i\boldsymbol{l}_i') \circ (\boldsymbol{M}_D\boldsymbol{M}_D') - (\boldsymbol{l}_i\boldsymbol{l}_i') \circ (\boldsymbol{l}_i\boldsymbol{l}_i')
\end{aligned} \tag{S15}$$

Using Eq. (S14) and (S15), we reduced the complexity of calculating $\boldsymbol{H}_i$ from $\mathcal{O}(n^2p^2)$ to $\mathcal{O}(n(n+r)^2p)$. Since $p$ is usually much larger than $n+r$, it is expected that $n(n+r)^2p$ is much smaller than $n^2p^2$.

### 5 Generalization of the model

As stated in the main text, the test of heterotic effects in our hQTL-ODS model is a kernel-based association test [6]. Thus, the framework can be used in more general context rather than for MPH only. Here, we mentioned two cases and both have already been implemented in hQTL-ODS as options.

#### 5.1 Testing the heterotic effect for better parent heterosis

The beter parent heterosis (BPH) is defined as the difference between the performance of the hybrid progeny and the superior parent, i.e., $y_{BPH,F} = y_F - \max\{y_{P1}, y_{P2}\}$. Let $\boldsymbol{y}_{ori}$ be the $(n+r)$-dimensional vector of the observed original trait values of all hybrids and parents, and $\boldsymbol{y}_{BPH}$ be the n-dimensional vector of the MPH values of all hybrids. Then, similar to the case of MPH, there is an $n \times (n+r)$ matrix of linear transformation $\boldsymbol{T}_b$ such that $\boldsymbol{y}_{BPH} = \boldsymbol{T}_b\boldsymbol{y}_{ori}$.

*Example.* Suppose that there are three parental lines $P_1$, $P_2$, $P_3$, and their three hybrid progenies $F_{1\times2}$, $F_{1\times3}$, $F_{2\times3}$. Order the entries of $\boldsymbol{y}_{ori}$ as $(\boldsymbol{y}_{P_1}, \boldsymbol{y}_{P_2}, \boldsymbol{y}_{P_3}, \boldsymbol{y}_{F_{1\times2}}, \boldsymbol{y}_{F_{1\times3}}, \boldsymbol{y}_{F_{2\times3}})'$. In addition, assume that $\boldsymbol{y}_{P_1} > \boldsymbol{y}_{P_2} > \boldsymbol{y}_{P_3}$. Then, we have

$$\boldsymbol{y}_{BPH} = \begin{bmatrix} y_{BPH,F_{1\times2}} \\ y_{BPH,F_{1\times3}} \\ y_{BPH,F_{2\times3}} \end{bmatrix} = \begin{bmatrix} -1 & 0 & 0 & 1 & 0 & 0 \\ -1 & 0 & 0 & 0 & 1 & 0 \\ 0 & -1 & 0 & 0 & 0 & 1 \end{bmatrix} \cdot \begin{bmatrix} \boldsymbol{y}_{P_1} \\ \boldsymbol{y}_{P_2} \\ \boldsymbol{y}_{P_3} \\ \boldsymbol{y}_{F_{1\times2}} \\ \boldsymbol{y}_{F_{1\times3}} \\ \boldsymbol{y}_{F_{2\times3}} \end{bmatrix}.$$

That is,

$$\boldsymbol{T}_b = \begin{bmatrix} -1 & 0 & 0 & 1 & 0 & 0 \\ -1 & 0 & 0 & 0 & 1 & 0 \\ 0 & -1 & 0 & 0 & 0 & 1 \end{bmatrix}.$$

If we consider MPH, we have

$$\boldsymbol{y}_{MPH} = \begin{bmatrix} y_{MPH,F_{1\times2}} \\ y_{MPH,F_{1\times3}} \\ y_{MPH,F_{2\times3}} \end{bmatrix} = \begin{bmatrix} -\frac{1}{2} & -\frac{1}{2} & 0 & 1 & 0 & 0 \\ -\frac{1}{2} & 0 & -\frac{1}{2} & 0 & 1 & 0 \\ 0 & -\frac{1}{2} & -\frac{1}{2} & 0 & 0 & 1 \end{bmatrix} \cdot \begin{bmatrix} \boldsymbol{y}_{P_1} \\ \boldsymbol{y}_{P_2} \\ \boldsymbol{y}_{P_3} \\ \boldsymbol{y}_{F_{1\times2}} \\ \boldsymbol{y}_{F_{1\times3}} \\ \boldsymbol{y}_{F_{2\times3}} \end{bmatrix},$$

which means that

$$\boldsymbol{T} = \begin{bmatrix} -\frac{1}{2} & -\frac{1}{2} & 0 & 1 & 0 & 0 \\ -\frac{1}{2} & 0 & -\frac{1}{2} & 0 & 1 & 0 \\ 0 & -\frac{1}{2} & -\frac{1}{2} & 0 & 0 & 1 \end{bmatrix}.$$

Similar to the heterotic effect for MPH (Eq. 4), we can define the heterotic effect of a QTL for BPH, denoted by $\boldsymbol{b}_i$:

$$\boldsymbol{b}_i = \boldsymbol{T}_b \left[ \boldsymbol{m}_i a_i + \boldsymbol{l}_i d_i + \frac{1}{2} \sum_{\substack{j=1 \\ j \neq i}}^{p} ((\boldsymbol{m}_i \circ \boldsymbol{m}_j))aa_{ij} + (\boldsymbol{m}_i \circ \boldsymbol{l}_j))ad_{ij} + (\boldsymbol{l}_i \circ \boldsymbol{m}_j))ad_{ji} + (\boldsymbol{l}_i \circ \boldsymbol{l}_j))dd_{ij} \right]. \quad \text{(S16)}$$

Note that the additive effects $a_i$ is included in the above equation, which is an important difference from Eq. (4) for MPH. The reason is that the additive effects do not cancel out for BPH, i.e. $\boldsymbol{T}_b \boldsymbol{m}_i \neq 0$ in general.

Then, we can use exactly the same approach to test $\boldsymbol{b}_i$. Namely, we start from the same model for the original trait as Eq. (5) in the Methods section:

$$\boldsymbol{y}_{ori} = \boldsymbol{1}_{n+r}\mu + \boldsymbol{X}_c\boldsymbol{\alpha} + \boldsymbol{M}_A\boldsymbol{a} + \boldsymbol{M}_D\boldsymbol{d} + \boldsymbol{M}_{AA}\boldsymbol{aa} + \boldsymbol{M}_{AD}\boldsymbol{ad} + \boldsymbol{M}_{DD}\boldsymbol{dd} + \boldsymbol{e}. \quad \text{(S17)}$$

Then, left-multiplying with $\boldsymbol{T}_b$, we obtain:

$$\boldsymbol{y}_{BPH} = \boldsymbol{T}_b\boldsymbol{1}_{n+r}\mu + \boldsymbol{T}_b\boldsymbol{X}_c\boldsymbol{\alpha} + \boldsymbol{T}_b\boldsymbol{M}_A\boldsymbol{a} + \boldsymbol{T}_b\boldsymbol{M}_D\boldsymbol{d} + \boldsymbol{T}_b\boldsymbol{M}_{AA}\boldsymbol{aa} + \boldsymbol{T}_b\boldsymbol{M}_{AD}\boldsymbol{ad} + \boldsymbol{T}_b\boldsymbol{M}_{DD}\boldsymbol{dd} + \boldsymbol{T}_b\boldsymbol{e}. \quad \text{(S18)}$$

Similar to the case of MPH, we have $\boldsymbol{T}_b\boldsymbol{1}_{n+r} = \boldsymbol{0}_n$. But in general $\boldsymbol{T}_b\boldsymbol{M}_A \neq \boldsymbol{0}_{n\times p}$. Let $\boldsymbol{X}_b = \boldsymbol{T}_b\boldsymbol{X}_c$, $\boldsymbol{g}_{A,b} = \boldsymbol{T}_b\boldsymbol{M}_A\boldsymbol{a}$, $\boldsymbol{g}_{D,b} = \boldsymbol{T}_b\boldsymbol{M}_D\boldsymbol{d}$, $\boldsymbol{g}_{AA,b} = \boldsymbol{T}_b\boldsymbol{M}_{AA}\boldsymbol{aa}$, $\boldsymbol{g}_{AD,b} = \boldsymbol{T}_b\boldsymbol{M}_{AD}\boldsymbol{ad}$, $\boldsymbol{g}_{DD,b} = \boldsymbol{T}_b\boldsymbol{M}_{DD}\boldsymbol{dd}$, and $\boldsymbol{\varepsilon}_b = \boldsymbol{T}_b\boldsymbol{e}$, where the subscript $b$ indicates BPH. Then, Eq. (S18) can be written as:

$$\boldsymbol{y}_{BPH} = \boldsymbol{X}_b\boldsymbol{\alpha} + \boldsymbol{g}_{A,b} + \boldsymbol{g}_{D,b} + \boldsymbol{g}_{AA,b} + \boldsymbol{g}_{AD,b} + \boldsymbol{g}_{DD,b} + \boldsymbol{\varepsilon}_b. \quad \text{(S19)}$$

This is the null model for testing the heterotic effects of BPH. In the model, $\boldsymbol{\alpha}$ is assumed to be a vector of fixed effects. Let $*$ denote $A$, $D$, $AA$, $AD$ or $DD$, and we assume that $\boldsymbol{g}_{*,b} \sim N(\boldsymbol{0}, K_{*,b}\sigma^2_{*,b})$, where $\boldsymbol{K}_{*,b} = \boldsymbol{T}_b\boldsymbol{M}_*\boldsymbol{M}'_*\boldsymbol{T}'_b/c_{*,b}$, $c_{*,b} = \frac{1}{n}\operatorname{tr}(\boldsymbol{T}_b\boldsymbol{M}_*\boldsymbol{M}'_*\boldsymbol{T}'_b)$. Moreover, $\boldsymbol{\varepsilon}_b \sim N(\boldsymbol{0}, \boldsymbol{T}_b\boldsymbol{T}'_b\sigma^2_{\varepsilon,b})$.

Adding $\boldsymbol{b}_i$ in Eq. (S19), we obtain the alternative model:

$$\boldsymbol{y}_{BPH} = \boldsymbol{X}_b\boldsymbol{\alpha} + \boldsymbol{b}_i + \boldsymbol{g}_{A,b} + \boldsymbol{g}_{D,b} + \boldsymbol{g}_{AA,b} + \boldsymbol{g}_{AD,b} + \boldsymbol{g}_{DD,b} + \boldsymbol{\varepsilon}_b, \tag{S20}$$

where $\boldsymbol{b}_i \sim N(\boldsymbol{0}, \boldsymbol{B}_i\sigma^2_{b_i})$, $\boldsymbol{B}_i = \boldsymbol{Z}_{i,b}\boldsymbol{Z}'_{i,b}/c_{i,b}$, $c_{i,b} = \mathrm{tr}(\boldsymbol{Z}_{i,b}\boldsymbol{Z}'_{i,b})$, $\boldsymbol{Z}_{i,b}$ is the $n \times (4p-2)$ matrix whose columns consist of $\boldsymbol{T}_b\boldsymbol{m}_i$, $\boldsymbol{T}_b\boldsymbol{l}_i$, $\frac{1}{2}\boldsymbol{T}_b(\boldsymbol{m}_i \circ \boldsymbol{m}_j)$, $\frac{1}{2}\boldsymbol{T}_b(\boldsymbol{m}_i \circ \boldsymbol{l}_j)$, $\frac{1}{2}\boldsymbol{T}_b(\boldsymbol{l}_i \circ \boldsymbol{m}_j)$ and $\frac{1}{2}\boldsymbol{T}_b(\boldsymbol{l}_i \circ \boldsymbol{l}_j)$ for $j = 1, 2, \ldots, p$ and $j \neq i$.

The significance of $\boldsymbol{b}_i$ is assessed by a likelihood ratio test comparing the null model (S19) and the alternative model (S20).

#### 5.2 Testing marginal epistatic effects for the original trait

Marginal epistatic effect refers to the combined pairwise interaction effects between a given genetic variant and all other variants in the genetic background [7]. If we consider the marginal epistatic effect of a locus for heterosis, it is actually very similar to the heterotic effect. But there is an important difference: the marginal epistatic effect does not contains any main (additive or dominance) effect. Here, we consider marginal epistatic effects for the original trait.

For any $i \in \{1, 2, \ldots, p\}$, the marginal epistatic effect of the $i$-th marker, denoted by $\boldsymbol{me}_i$, is defined as:

$$\boldsymbol{me}_i = \sum_{\substack{j=1 \\ j\neq i}} \big((\boldsymbol{m}_i \circ \boldsymbol{m}_j)aa_{ij} + (\boldsymbol{m}_i \circ \boldsymbol{l}_j)ad_{ij} + (\boldsymbol{l}_i \circ \boldsymbol{m}_j)ad_{ji} + (\boldsymbol{l}_i \circ \boldsymbol{l}_j)dd_{ij}\big) \tag{S21}$$

Again, we can use exactly the same approach to test $\boldsymbol{me}_i$ as developed for $\boldsymbol{h}_i$ in the main text. In fact, the models are simpler because we do not have to apply any linear transformations. More precisely, we take Eq. (S17) as the null model. Let $\boldsymbol{g}_{A,o} = \boldsymbol{M}_A\boldsymbol{a}$, $\boldsymbol{g}_{D,o} = \boldsymbol{M}_D\boldsymbol{d}$, $\boldsymbol{g}_{AA,o} = \boldsymbol{M}_{AA}\boldsymbol{aa}$, $\boldsymbol{g}_{AD,o} = \boldsymbol{M}_{AD}\boldsymbol{ad}$ and $\boldsymbol{g}_{DD,o} = \boldsymbol{M}_{DD}\boldsymbol{dd}$, where the subscript $o$ indicates the original trait. Then, (S17) can be rewritten as:

$$\boldsymbol{y}_{ori} = \boldsymbol{1}_{n+r}\mu + \boldsymbol{X}_c\boldsymbol{\alpha} + \boldsymbol{g}_{A,o} + \boldsymbol{g}_{D,o} + \boldsymbol{g}_{AA,o} + \boldsymbol{g}_{AD,o} + \boldsymbol{g}_{DD,o} + \boldsymbol{e}. \tag{S22}$$

In the model, $\mu$ and $\boldsymbol{\alpha}$ are assumed to be fixed effects. Let $*$ denote $A$, $D$, $AA$, $AD$ or $DD$, and we assume that $\boldsymbol{g}_{*,o} \sim N(\boldsymbol{0}, K_{*,o}\sigma^2_{*,o})$, where $\boldsymbol{K}_{*,o} = \boldsymbol{M}_*\boldsymbol{M}'_*/c_{*,o}$, $c_{*,o} = \frac{1}{n+r}\,\mathrm{tr}(\boldsymbol{M}_*\boldsymbol{M}'_*)$, and $\boldsymbol{e} \sim N(\boldsymbol{0}, \boldsymbol{I}_{n+r}\sigma^2_e)$.

Adding $\boldsymbol{me}_i$ in (S22), we obtain the alternative model:

$$\boldsymbol{y}_{ori} = \boldsymbol{1}_{n+r}\mu + \boldsymbol{X}_c\boldsymbol{\alpha} + \boldsymbol{me}_i + \boldsymbol{g}_{A,o} + \boldsymbol{g}_{D,o} + \boldsymbol{g}_{AA,o} + \boldsymbol{g}_{AD,o} + \boldsymbol{g}_{DD,o} + \boldsymbol{e}, \tag{S23}$$

where $\boldsymbol{me}_i \sim N(\boldsymbol{0}, \boldsymbol{E}_i\sigma^2_{me_i})$, $\boldsymbol{E}_i = \boldsymbol{Z}_{i,me}\boldsymbol{Z}'_{i,me}/c_{i,me}$, $c_{i,me} = \mathrm{tr}(\boldsymbol{Z}_{i,me}\boldsymbol{Z}'_{i,me})$, $\boldsymbol{Z}_{i,me}$ is the $n \times (4p-4)$ matrix whose columns consist of $\boldsymbol{m}_i \circ \boldsymbol{m}_j$, $\boldsymbol{m}_i \circ \boldsymbol{l}_j$, $\boldsymbol{l}_i \circ \boldsymbol{m}_j$ and $\boldsymbol{l}_i \circ \boldsymbol{l}_j$ for $j = 1, 2, \ldots, p$ and $j \neq i$.

The significance of $\boldsymbol{me}_i$ is assessed by a likelihood ratio test comparing the null model (S22) and the alternative model (S23).

In certain cases, we may only consider the marginal additive-by-additive epistatic effects (as in [7]) or it is only possible to do so (e.g., in a inbred population). Then, the definition (S21) is simplified as

$$\boldsymbol{me}_i = \sum_{\substack{j=1 \\ j\neq i}} (\boldsymbol{m}_i \circ \boldsymbol{m}_j)aa_{ij} \tag{S24}$$

The null model (S22) and the alternative model (S23) are simplified as:

$$\begin{aligned} \boldsymbol{y}_{ori} &= \boldsymbol{1}_{n+r}\mu + \boldsymbol{X}_c\boldsymbol{\alpha} + \boldsymbol{g}_{A,o} + \boldsymbol{g}_{AA,o} + \boldsymbol{e}. \\ \boldsymbol{y}_{ori} &= \boldsymbol{1}_{n+r}\mu + \boldsymbol{X}_c\boldsymbol{\alpha} + \boldsymbol{me}_i + \boldsymbol{g}_{A,o} + \boldsymbol{g}_{AA,o} + \boldsymbol{e}. \end{aligned} \tag{S25}$$

where the model assumptions are the same as for (S22) and (S23), except that $\boldsymbol{Z}_{i,me}$ is now the $n \times (p-1)$ matrix whose columns consist of $\boldsymbol{m}_i \circ \boldsymbol{m}_j$ for $j = 1, 2, \ldots, p$ and $j \neq i$.
